# Transcription factor activity profiling reveals the role of REST and LEF1 in the recovery from depression

**DOI:** 10.1101/2023.11.30.569491

**Authors:** Hajime Yamamoto, Satomi Araki, Ryoma Onodera, Yasuhiro Go, Kentaro Abe

## Abstract

Psychophysiological disorders chronically impair brain functions, often accompanied by dysregulation of multiple genes, suggesting a multifaceted etiology behind the symptoms. To explore transcription factors (TFs) involved in such transcriptomic changes, we analyzed TF-activity profiles (TFAPs) from the brains of mice experienced chronic stress, and revealed alteration in TF-activity correlating with their pathophysiological phenotypes. We identified REST/NRSF and TCF/LEF associated with depressive phenotypes and discovered that neuropsychiatric drugs sertraline and lithium influence REST- and TCF/LEF-activity, both in vitro and in vivo, thereby affecting gene expression profiles. Pharmacological or genetic manipulation of REST- or TCF/LEF-activity in defeated mice impacts post-stress recovery from depressive phenotypes, with combined treatment further augmenting the outcomes. Our TFAP analysis enhances understanding of molecular mechanisms underpinning chronic diseases, aiding future therapeutic strategy development.

## Introduction

Depression is one of the most prevalent mental disorders characterized by chronic brain dysfunction, causing symptoms like depressed mood, loss of pleasure, and fatigue (*1*, *2*). Our current understanding of the molecular mechanism underlying depression remains insufficient to establish effective remedies for all patients suffering from the disease (*3*, *4*). One of the most influential hypotheses about the molecular cause of depression is the monoamine hypothesis (*5–7*). Although drugs that modulate the synaptic concentration of monoamine serotonin are the main focus of current medication against depression, their causal role in depression remains a subject of debate due to many observations that do not conform to the hypothesis (*7*, *8*). Notably, the therapeutic effect of antidepressants targeting monoamines often manifests only after weeks of treatment (*9*), implying that their effectiveness requires chronic changes in the brain. Studies on diverse animal models have revealed transcriptomic changes in the brains of depressed animals (*10*, *11*), and the post-mortal brain sample of human patients confirms this (*12*, *13*). These observations suggest that plastic changes in the brain are involved in both the pathogenesis of the disease and its therapeutic response. Although several genes have been suggested to play a role in the development of the disease, our complete understanding of the pathophysiology of depression is still in its infancy, due to the involvement of multiple genes and their interaction with environmental risk factors (*2*, *10*, *14–16*).

Considering the critical function of coordinated modulation in the expression and functions of numerous genes during the initiation and the recuperation phase of chronic neural disorders, such as depression, identifying the key molecules that orchestrate this complex regulatory mechanism is crucial. Transcription factors (TFs) emerge as potential candidates for these core molecules, owing to their capacity to interact with multiple gene regulatory domains, thereby controlling the expression of target gene sets. Given this characteristic, it is plausible that TF malfunction plays a role in numerous diseases (*17–19*). Moreover, investigating TF-activities offers advantages over transcriptome analysis, since manipulating TFs facilitates practical intervention in systems that intricately involve multiple genes. Therefore, elucidating the activities of multiple TFs in chronic diseases, including depression, can contribute to understanding their pathophysiology and aid in the development of therapeutic strategies against them. Recently, we have developed a method to quantitatively analyze the activity of multiple TFs using a viral-vector-based TF-activity-reporter battery (*20*, *21*). This method enable us to obtain the TF-activity profile (TFAP) from a specific brain population *in vivo*. Additionally, it allows for the comparison of TF-activity signatures across mice with distinct behavioral or pathophysiological conditions. In this study, we applied this method to investigate changes in TF-activity of the depression model mice, particularly focusing on the prefrontal cortex (PFC), a region consistently impaired and playing a crucial role in establishing depression symptoms and recovery (*22*, *23*).

## Results

### Transcription factor activity profiling of stress-induced depression

To get a molecular insight into the chronic change of the brain in pathological conditions, we obtained the TFAP of mice who experienced chronic social defeat stress, a well-established rodent model of depression (*4*). In concrete, we injected the lentivirus into the ventricle of the embryo *in utero* to express the TF-activity-reporters in a defined cell population after they were aged > 8 weeks (Fig. 1A) (*24*). Mice expressing TF-activity-reporter constructs were subjected to repeated social defeat stress for 10 days (Fig. 1B), which resulted in a tendency for depressed behaviors such as social avoidance and anxiety-like behavior (Figs. 1C–D and Supplementary Figs. S1A–C). Social interaction test revealed the defeated mice showed a chronic reduction of social interaction rate (SI-rate), which gradually returned to that of control mice by 15 days (Supplementary Fig S1A). The defeated mice were grouped into resilient and susceptible groups according to their SI-rate (Fig. 1D) (*25*). We measured the activities of 30 TF-reporters in the anterior portion of the cortex in the defeated mice 3 days after the stress period ended. These measurements were corrected for the mean value in the control group and then integrated as a TFAP (Fig. 1E, see Methods). This analysis indicates that TF-activities *in vivo* were chronically affected by repeated exposure to stress. The shapes of TFAPs from resilient and susceptible were distinct from each other, with a total Euclidean distance of 16.4. Simulation-based analysis on the change in TF-activity revealed that the TFAP of susceptible mice demonstrated the most pronounced characteristics with an increased variation of TF-activities (Supplementary Figs. S2A, C, D; see Methods).

**Fig. 1.**
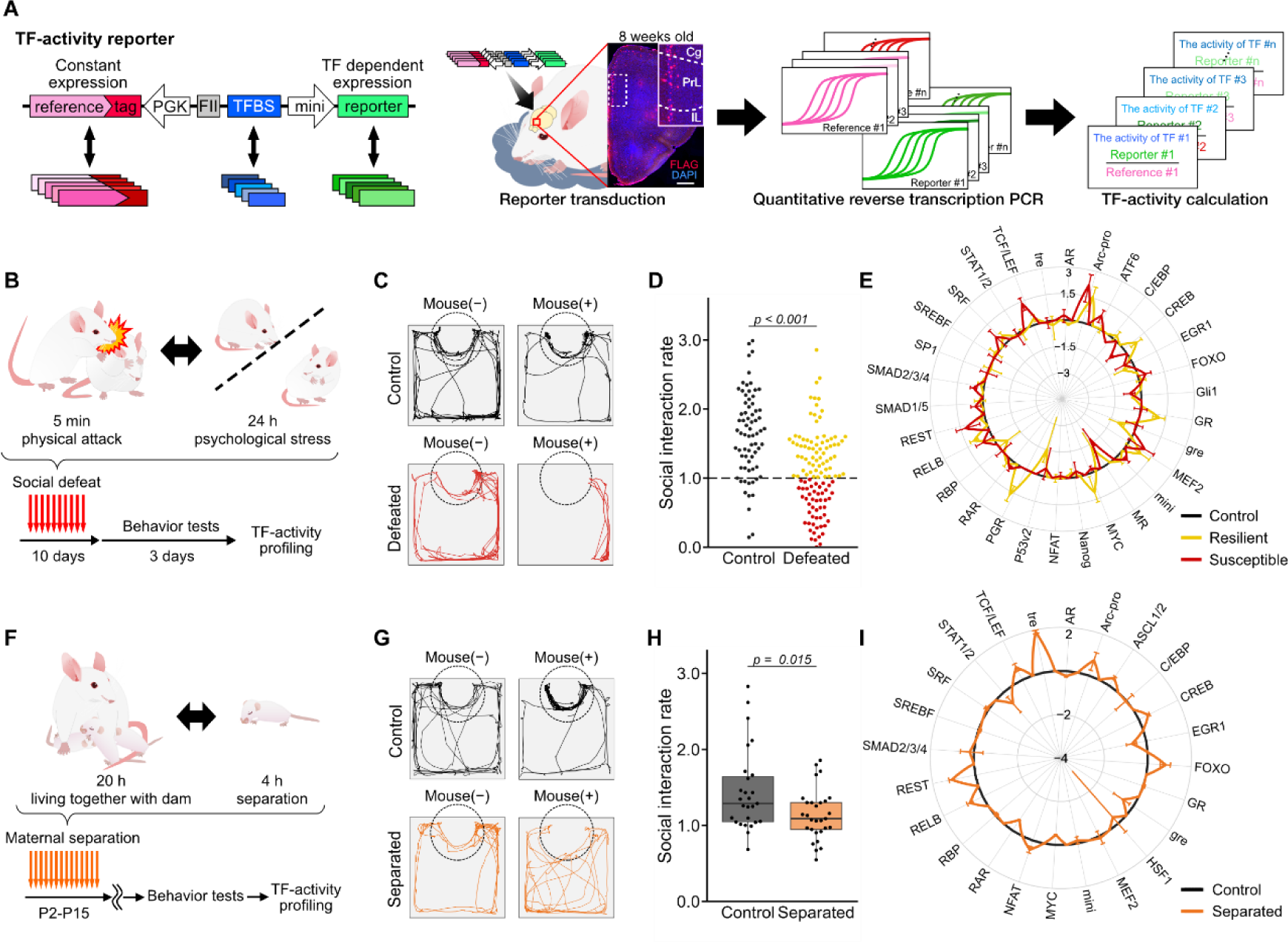
TF-activity profile of stressed mice. (**A**) Schematic illustration of TF-activity profiling *in vivo*. The brain image shows the reporter infected cells (red, flag-tag) in the prefrontal cortex. (**B–E**) Effects of chronic social defeat stress. (**B**) Scheme of the repeated social defeat and TF-activity profiling. (**C**) Trajectories in social interaction test. The dotted line shows the interaction zone where an experimental mouse can interact with an unfamiliar mouse. (**D**) SI-rates of control (black), resilient (yellow), and susceptible (red), Welch’s *t*-test, *n* = 71, 119. (**E**) Radar graph showing changes in TF-activities by social defeat stress. For each TF, corrected mean ± sem of TF-activity are shown as log_2_-fold-changes. See Methods for details. (**F–I**) Effects of maternal separation stress. (**F**) Scheme of the maternal separation and TF-activity profiling. (**G**) same as (**C**). (**H**) SI-rates of control (black) and separated mice (orange) at 8-week-old. Welch’s *t*-test; *n* = 27, 29. (**I**) Same as (**E**), showing changes in the activity of TFs by maternally separated stress.

Several experimental models in rodents have been reported to show similar depressive phenotypes (*26*, *27*). Early-life maternal separation is one such animal model, resulting in depressiveness and sensitization to stress after growing (*28*). We examined a TFAP of the mice who experienced maternal separation in their early life. In concrete, mice expressing TF-activity-reporter constructs were subjected to repeated maternal separation stress for 14 days at postnatal day 2–15, after which they were raised under the same condition as the non-separated siblings till eight weeks post-birth (Fig. 1F). Those mice at eight weeks age demonstrated a reduced tendency towards social interaction (Figs. 1G–H and Supplementary Figs. S1D–E), a phenotype reminiscent of mice subjected to social defeat stress. The shape of TFAP from the adult mice that experienced maternal separation was distinct from that of the control, with a total Euclidean distance of 11.4 (Fig. 1I). Simulation-based analysis on change of TF-activity revealed that TFAP in maternally separated mice became highly diverse (Supplementary Figs. S2B, E), indicating the activities of multiple TFs was chronically affected by experiencing early-life stress. These observations suggest that the endogenous activities of TFs undergo chronic alterations in the mice showing depressive phenotypes, and the degree of change in TFAPs is related to the observed behavioral phenotype.

### TFAP analysis reveals the role of potential TFs involved in stress responses

The stressed experience should impress a distinctive signature on the activity profile of TFs. We hypothesized that the shared phenotype observed in two stress models arose from a combination of TFs exhibiting consistent activity under both conditions. To identify the potential combination of TFs responsible for producing the phenotype, we conducted a multivariate cosine similarity analysis of the TF-activities. In this analysis, we created vectors whose components representing the value of each vertex on the TFAP for a variety of TF combinations. Subsequently, for every combination of 3–6 TFs, we plotted the cosine similarity along with their product between the vectors of the two stress conditions (Fig. 2A, see Methods). This process enriches plots representing the largest vector magnitude and highest cosine similarity to the up-right portion (Supplementary Fig S3A). We discovered that vectors reflecting three TF-activity-reporters are significantly enriched at the up-right portion, indicating that those TFs are the primary candidate for the TF combination that correlates with the shared depressive phenotype between the two stress conditions (Supplementary Figs. S3B–F). Those three were found to be the reporters reflecting the activity of TCF/LEF (T cell factor / lymphoid enhancer factor family), REST (RE1 silencing transcription factor / NRSF), and endogenous Arc/Arg3.1 promoter (*29*) (Fig. 2B).

**Fig. 2.**
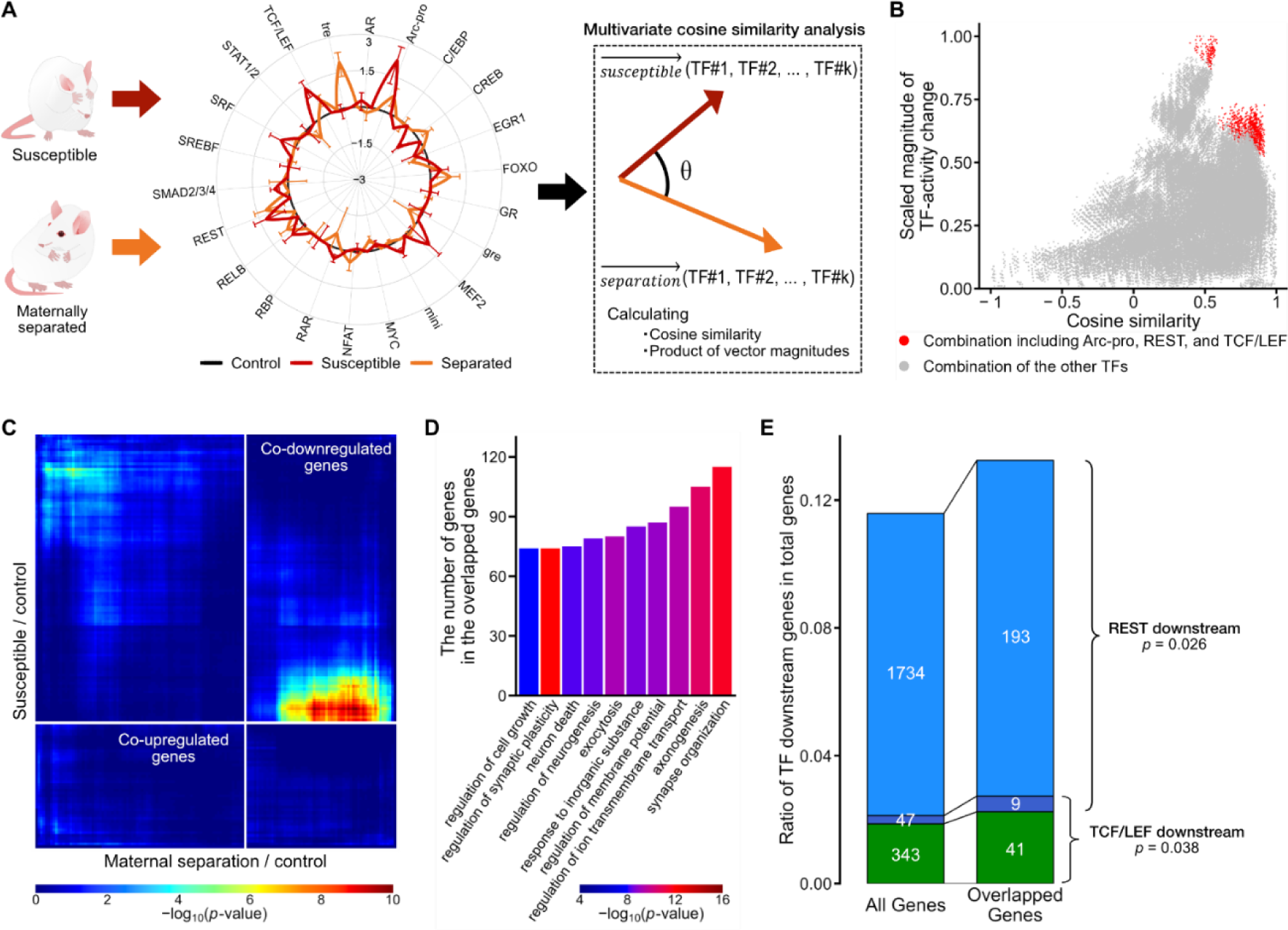
Multi-omics identification of TFs involved in depressive phenotype. (**A**) Conceptual diagram of multivariate cosine similarity analysis for identifying TFs with similar activity change. Vectors consisting of *k* components, selected from the TFAPs of the susceptible and maternally separated mice were analyzed. See Supplementary Fig. S3 for details. (**B**) The scatter plot shows the result of the analysis against every 3 to 6 combinations of TFs. TF combinations that include Arc-pro, REST, and TCF/LEF are shown in red. (**C**) The RRHO map of transcriptome changes in the two stressed conditions (social defeat and maternal separation). Pixels in the heat map represent the degree of significance of overlap. (**D**) GO analysis of the significantly downregulated 1845 genes identified by the RRHO analysis. The top 10 GO terms in “biological process” are shown. The color represents adjusted *p*-values, which are calculated by hypergeometric distribution, using the Benjamini-Hochberg procedure. (**E**) The cumulative bars indicate the ratio and number of genes downstream of REST, TCF/LEF in the RNA-seq analysis (All genes, 21355 genes; overlapped genes, 1916 genes identified by RRHO analysis). The *p*-values were calculated based on a hypergeometric probability distribution.

What influence did the observed change in TF-activity have on the depressive phenotype? To unravel this, we conducted transcriptome analysis using RNA-seq on the mouse brains from which the TFAP was obtained. Between the control group and susceptible mice, we identified 174 upregulated genes (> 2 fold) and 319 downregulated genes (< 0.5 fold) in their anterior cortex. Similarly, between the control and maternally separated mice, we detected 405 upregulated (> 2 fold) and 242 downregulated genes (< 0.5 fold). A Rank-Rank Hypergeometric Overlap (RRHO) analysis reveals a set of genes exhibiting common expression patterns across two independent experimental conditions (*30*). Using the RRHO analysis for the transcriptomic data from the two depressive mice, i.e., the susceptible mice in the social defeat model and mice experienced early-life maternal separation stress, we identified 71 genes with upregulated and 1845 genes with downregulated in common (Fig. 2C). Gene ontology (GO) analysis indicated that this downregulated gene set was associated with neuronal functions (Fig. 2D), indicating that the brain function was impaired in the depressed mice through alterations the expression of this particular gene set. Promoter analysis of those overlapped gene sets revealed a significant enrichment of the regulon for both REST and TCF/LEF (Fig. 2E). Collectively, our multi-omics analysis of two depressive models revealed the potential involvement of specific TFs, TCF/LEF and REST, in the depressive phenotype in mice through the regulation of their downstream genes.

### Role of TCF/LEF in depression

Next, from the TFs whose activity was changed in the depressed mouse, we focused on the role of TCF/LEF, whose activity was significantly increased in the susceptible mice (Dunnett’s test; *p* = 9.82 × 10^−3^ versus control, *n* = 14 and 10; Fig. 1E). TCF/LEF is known to play a role in Wnt signaling pathway, which is involved in processes such as the proliferation or differentiation of stem cells (*31*, *32*). However, its role in the adult brain is not well established. Immunostaining of brain sections revealed that *Lef1*, one of the TFs in the TCF/LEF family, was expressed in the neurons of the cortex in susceptible mice (Fig. 3A). Consequently, we assessed whether the upregulation of TCF/LEF signaling influences the depressive phenotype in mice.

**Fig. 3.**
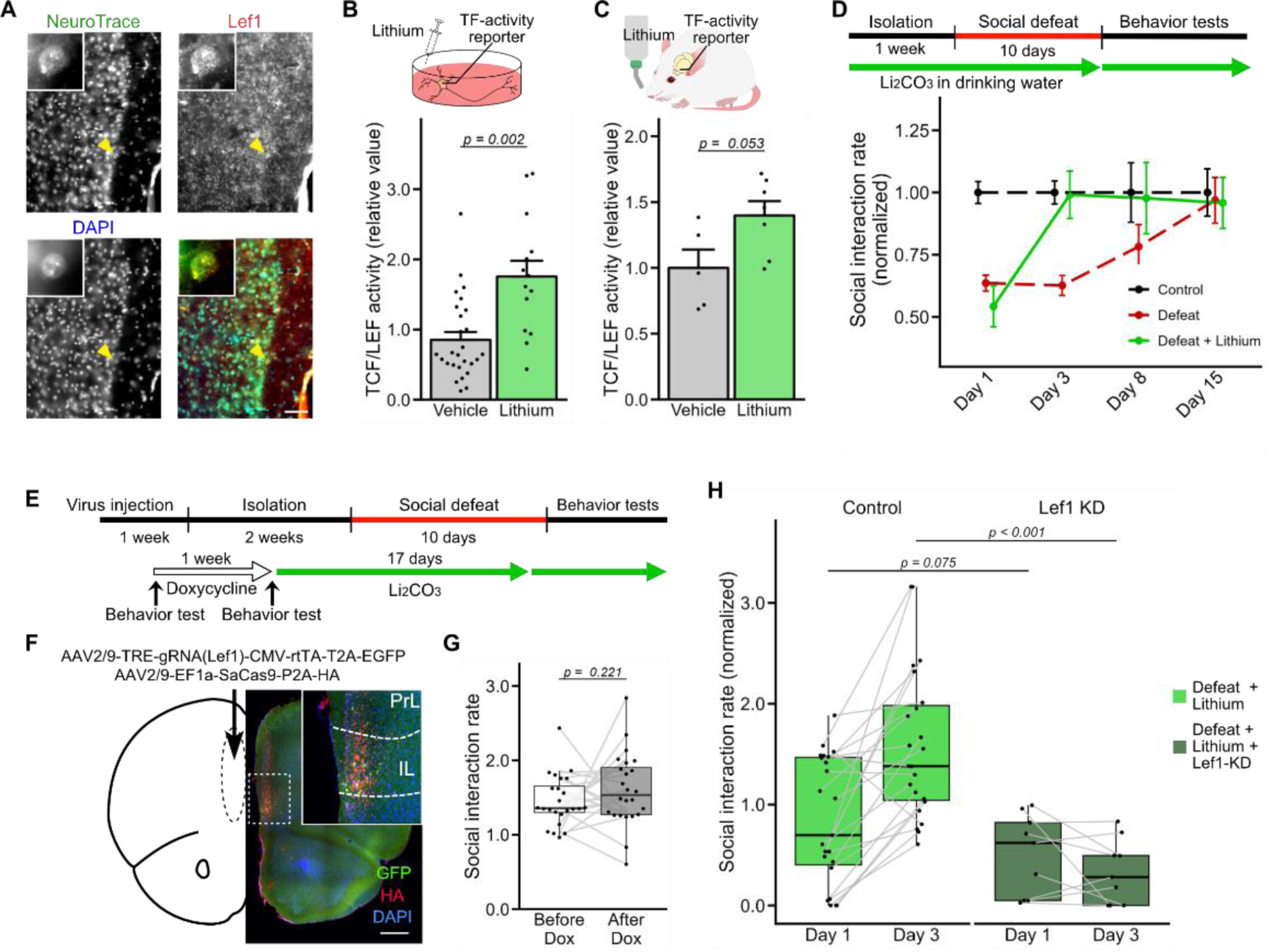
Role of TCF/LEF in the depressive phenotype. (**A**) An immunostained section of the anterior cortex. Scale bar; 100 mm. (**B–C**) TCF/LEF activity of cortical culture and the anterior cortex after lithium treatment. Mean ± sem; Welch’s *t*-test; vehicle, 27; lithium, 14 cultures in (**B**); vehicle, 5; lithium, 7 mice in (**C**). (**D**) SI-rate change after defeat stress with lithium treatment. Experimental schedule (top). SI-rates, normalized to control group on each day (bottom). The data include those shown in Fig. 1. Control, defeat, and defeat + lithium: day 1, *n* = 83, 126, 23; day 3, *n* = 83, 126, 23; day 8, *n* = 25, 34, 19; day 15, *n* = 22, 29, 17. (**E–H**) Social behaviors of *Lef1* knockdowned (Lef1-KD) mice. Experimental schedule (**E**), and an example of the brain section (**F**). Scale bar; 500 mm. (**G**) SI-rates before and after doxycycline treatment without stress. Paired *t*-test, 24 mice. (**H**) SI-rates at day 1 and day 3 after the 10-day social defeat, normalized to the defeat group in defeat + lithium mice, and to defeat + sham operated group in defeat + lithium + Lef1-KD mice. Welch’s *t*-test; defeat + lithium, 23; defeat + lithium + Lef1-KD, 8 mice in both post-stress days.

Lithium, a drug prescribed as a mood stabilizer, is known to inhibit GSK3β, thereby activating TCF/LEF activity (*33*). As expected, we observed an upregulation of TCF/LEF activity in cultured cortical neurons treated with lithium (20 mM for 24 h; Fig. 3B) and in the brain of mice with chronic administration of lithium (100 mg/kg/day for 17 days; Fig. 3C). We observed that lithium treatment alone did not affect social interaction tendency and behaviors in the open field (Supplementary Figs. S4A–C), suggesting that the upregulation of TCF/LEF activity does not induce a depressive phenotype. Next, we assessed the effects of lithium treatment on the mice that experienced chronic social defeat stress. After a 10-day social defeat stress, lithium-treated mice exhibited depressive phenotypes similar to vehicle-treated mice with no significant effect on SI-rate and open field test, indicating that lithium did not prevent the establishment of depression following the stressed experience (Fig. 3D and Supplementary Figs. S4D–E). Intriguingly, however, their SI-rate on the third day of post-stress was significantly increased and restored to a level comparable to that of non-stressed control mice (Fig. 3D). The effect size, quantified using Hedge’s *g* on the three days after the defeat stress, was 0.81, indicating a potent antidepressant effect of lithium. To investigate the role of TCF/LEF in this lithium-dependent post-stress recovery, we performed a doxycycline-dependent CRISPR/Cas9 mediated knockdown of endogenous Lef1 expression in the PFC (Figs. 3E–F and Supplementary Fig. S5). The knockdown of the endogenous *Lef1* did not affect their behaviors before the stress exposure (Fig. 3G). However, after being subjected to chronic stress, *Lef1* knockdown suppressed the effect of lithium to stimulate the post-stress recovery of the SI-rate (Fig. 3H). Together, these observations reveal a potential role of the TCF/LEF family in stressed mice: increased TCF/LEF activity does not affect the development of the depressive phenotype but promotes recovery from depression.

### Role of REST in depression

Next, we examined the role of REST, a transcription repressor, whose roles are well studied in the field of neural development (*34*, *35*), but its functions in mature neurons are also recognized (*36*). A recent study has indicated that sertraline, a widely prescribed antidepressant, interacts with mSin3, a co-factor of REST, to affect its function (*37*). Therefore, we assessed the impact of sertraline on REST-mediated gene regulation and mice behavior. Sertraline treatment (5 mg/kg/day for 17 days) significantly suppressed the decline in sociality of socially defeated mice (Fig. 4A). The effect sizes, quantified using Hedge’s *g*, were 0.75, 0.18, 0.36, and 0.45 at 1, 3, 8, and 15 days post social defeat, respectively. Sertraline treatment also affected anxiety-related behaviors after stress, as evidenced by increased time spent in the center zone, confirming its effectiveness as an antidepressant (Supplementary Figs. S6A–B). TF-activity-reporter analysis revealed that after administrating sertraline for 17 days, an upregulation of REST-activity was observed (Figs. 4B–C). On the other hand, chronic administration of sertraline did not affect the social tendency of non-stressed mice (Fig. 4D and Supplementary Figs. S6C–D), indicating that one of the mechanisms of sertraline for its antidepressant-effect may be by affecting REST-dependent gene regulation *in vivo* in the stressed individuals. To investigate this, we analyzed the transcriptome in the anterior region of the cortex in defeated mice and found significant changes in the depression-induced expression of REST regulons following treatment with sertraline, compared to the non-target genes (Fig. 4E). We observed that the sertraline exhibited a tendency to increase the expression of REST-regulons following defeat stress, compared to subjects treated with a control vehicle (Fig. 4F). These findings suggest that sertraline, in association with its established antidepressant properties, modulates gene expression *in vivo* through the modulation of REST-activity.

**Fig. 4.**
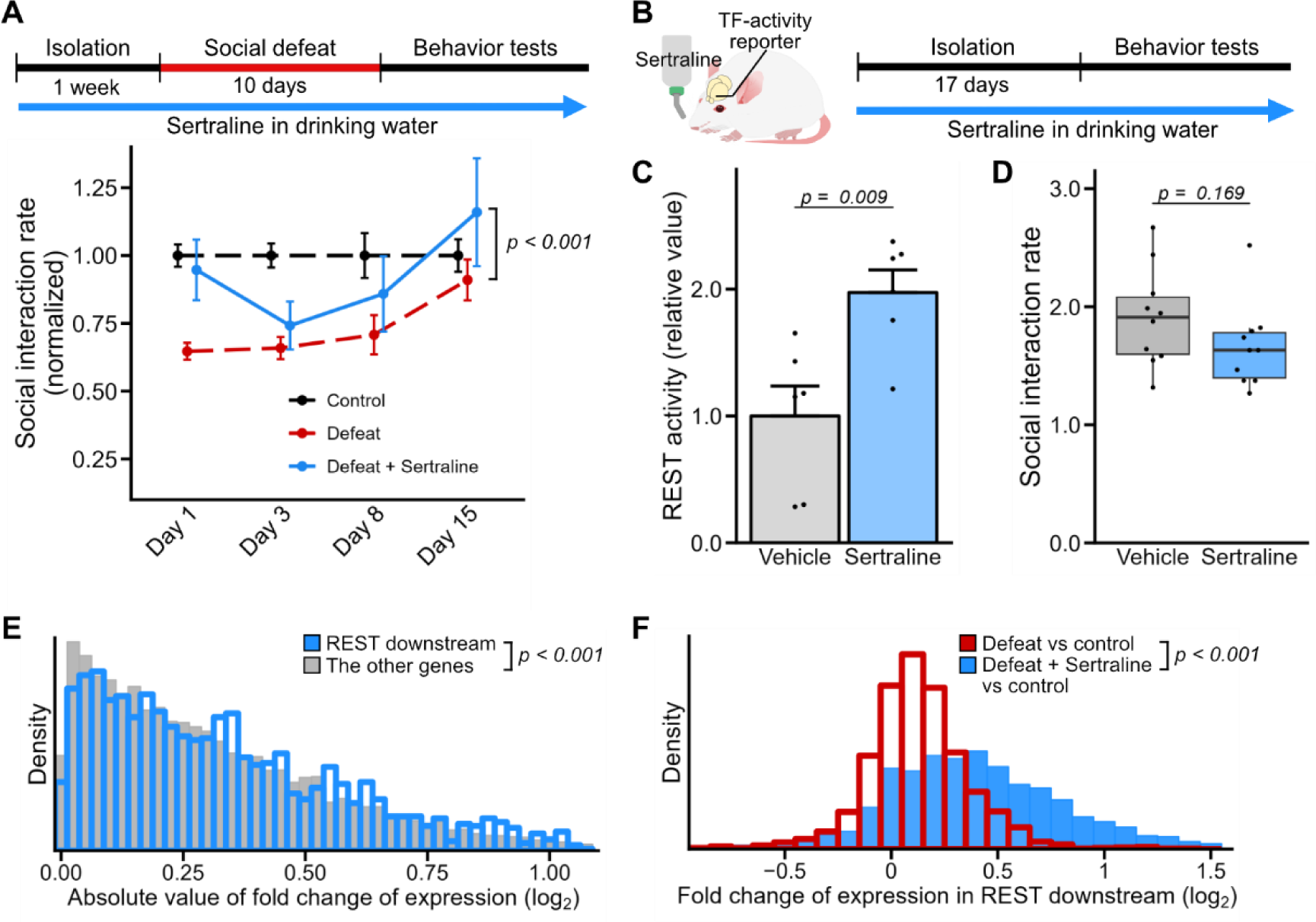
Effect of sertraline on REST-mediated gene transcription *in vivo*. (**A**) Experimental schedule of *in vivo* sertraline treatment (top) and SI-rate change after 10 days of social defeat with sertraline treatment (bottom). The data include those shown in Fig. 1. SI-rates were normalized to control group on each day. Two-way ANOVA in defeat and defeat + sertraline. The number of mice for control, defeat, and defeat + sertraline: day 1, 90, 126, 21; day 3, 90, 126, 21; day 8, 27, 38, 15; day 15, 24, 33, 15. (**B**) Experimental schedule of sertraline treatment to non-stressed mice. (**C**) Effect of sertraline on REST-activity in the anterior region of cortex. Welch’s *t*-test; vehicle, 6; sertraline, 6 mice. (**D**) SI-rates of vehicle-(grey) or sertraline-treated (light blue) mice without social defeat (Welch’s *t*-test; control, 10; sertraline, 10 mice). (**E**) Histogram of fold-change differences of gene expressions in the anterior cortex of defeated mice with sertraline treatment against vehicle-treated susceptible mice. Wilcoxon signed-rank test; REST downstream, 1477; others, 17009 genes. (**F**) Fold-change of expression level in REST downstream genes; red, control versus defeat; blue, control versus defeat + sertraline. Paired *t*-test, 1460 genes.

The phenotype described above could be attributable to well-established molecular mechanisms of sertraline as a selective serotonin reuptake inhibitor (SSRI), which is believed to increase serotonin’s synaptic concentration. To further investigate whether sertraline affects gene expression through modulating the function of REST, we evaluated the impact of sertraline on cortical neuron culture that lacks serotonergic neurons (Supplementary Figs. S7A–C) (*38*). In cortical culture *in vitro*, sertraline treatment upregulated the reporter activity of REST, similarly as observed *in vivo* (20 mM for 16 h; Supplementary Figs. S7D–E). Transcriptome analysis revealed an alteration in gene expression between vehicle- and sertraline-treated cortical neurons, with REST-regulons being significantly affected (Supplementary Figs. S7G–H). Expressing REST4, a splicing isoform of REST that is shown to affect the function of REST (*39*), reduced the sertraline-dependent REST-activity change (Supplementary Fig. S7F) and increased the kurtosis of the distribution of the expression-change of REST-regulons (Supplementary Figs. S7I–J). These suggest that changes in gene expression associated with REST activity were suppressed. Together, these results indicate that sertraline alters the transcriptomic profile in neurons by affecting the activity of REST. The widely acknowledged antidepressant effect of sertraline may be induced not only by its role as an SSRI but also through its impact on REST-mediated gene expression.

### Cooperative role of TCF/LEF and REST in the recovery from depression

Our TFAP analysis identified the involvement of TFs, TCF/LEF and REST, as well as the neural activity reporter Arc/Arg3.1, in mice exhibiting depressive phenotype (Fig. 2). Given that the drugs affecting either TCF/LEF or REST impacted the phenotype, we proceeded to examine the synergistic effect of TCF/LEF and REST on depression. For this purpose, we simultaneously measured the activity of TCF/LEF and REST in a single mouse subjected to social defeat stress. We noticed that the activities of both TCF/LEF and REST tended to exhibit an anti-parallel correlation with their SI-rate in individual mice, in both the control and defeated groups (Fig. 5A). Moreover, the correlation coefficient between the SI-rate in defeated mice was lower when the activities of TCF/LEF and REST were analyzed jointly, compared to when each TF was analyzed singly (Fig. 5B). This indicates that both TCF/LEF and REST play a role in the depressive phenotype. To further reveal the involvement of both TCF/LEF- and REST-activities at the cellular level, we performed TF-activity measurements and transcriptome analysis on single neurons using single nucleus RNA-seq (see Methods). We observed that the activity of both TCF/LEF and REST were highly heterogeneous among cells, and the heightened activity induced by defeat stress was distributed broadly, not confined to specific cell populations (Figs. 5C–E and Supplementary Fig. S8). Furthermore, transcriptome analysis of the defeated mice treated with either lithium or sertraline alone revealed that the alterations in gene expression induced by each drug were markedly different (Figs. 5F–G). These observations suggest that both TCF/LEF and REST contribute to changes in cell physiology, leading to an anti-depressive phenotype.

**Fig. 5.**
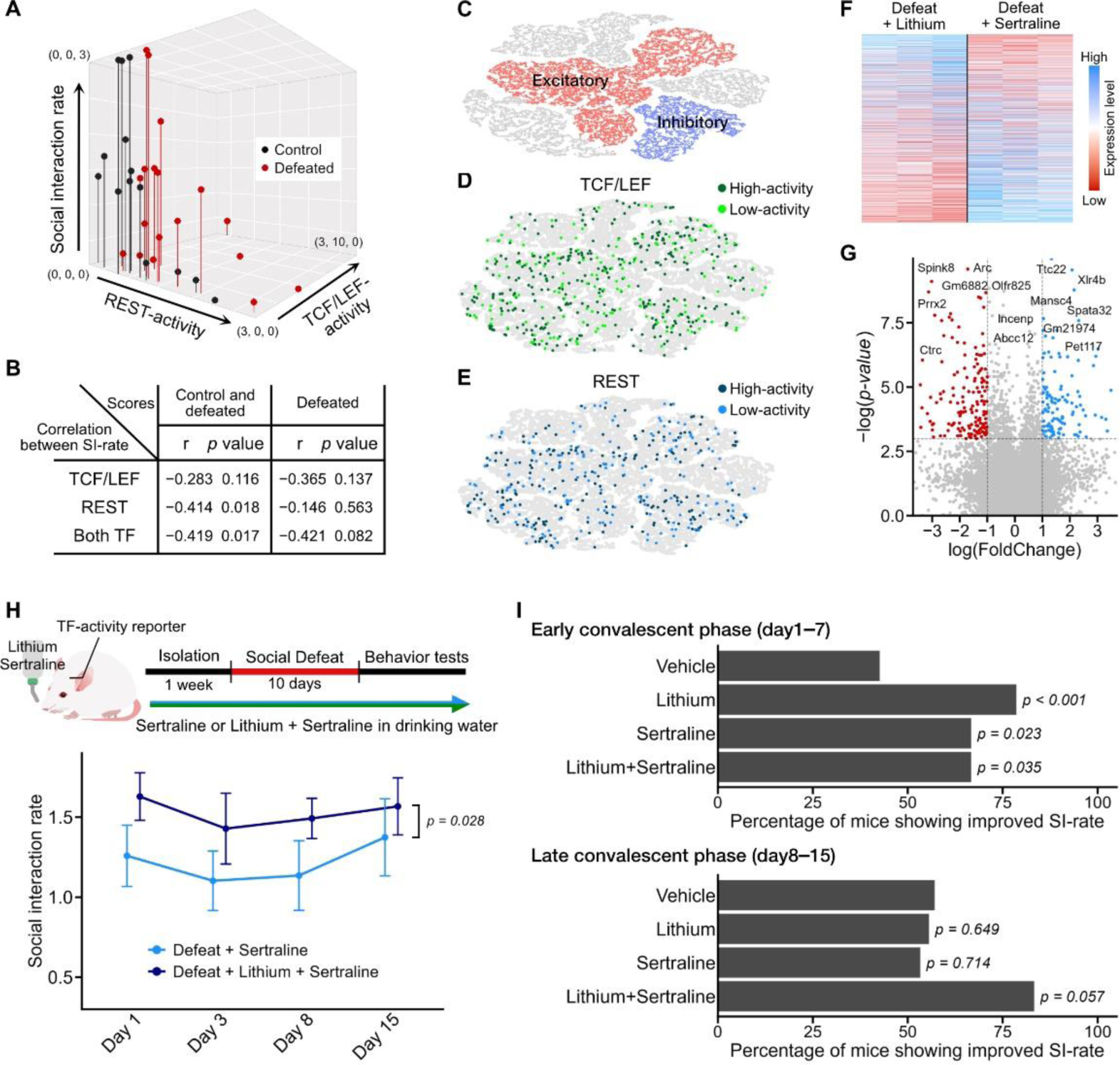
Individual differences in the activity of TCF/LEF and REST. (**A–B**) The SI-rate and the TF-activities for each mouse expressing both TCF/LEF- and REST-reporters. 3D plot (**A**) and the Pearson correlation analysis (**B**). Control, 14; defeat, 18 mice. (**C–E**) TF-activity analysis on single cell level in the PFC neurons. (**C**) The t-SNE plot illustrates the clustered cells after social defeat. See fig. S8 for details. (**D–E**) Distribution of TF-activity for TCF/LEF (**D**) and REST (**E**). Each TF-activity was compared to the control average within the clusters, and cells with high-activity and low-activity are indicated in dark/light color. (**F–G**) Bulk-RNA-seq of the anterior cortex of the defeated mice treated with lithium or sertraline (*n* = 3 each). The gene expression level (**F**), and the fold-change among two drug treatments (20565 genes) (**G**). The dotted lines show *p*-value of 0.05 and a log_2_-fold-change of ±1. (**H**) Effect of multiple drug treatment. Experimental scheme (top) and SI-rates change. Two-way ANOVA; the number of mice in each cohort: sertraline and lithium + sertraline; day 1, 14, 12; day 3, 15, 12; day 8, 15, 12; day 15, 15, 12. (**I**) Changes in depressive symptoms in mice after drug treatment during days 1–7 (top) and days 8–15 (bottom). The *p*-values were calculated based on a binomial test compared to vehicle-treated conditions.

In human patients with depression, multiple antidepressants are occasionally used in combination when the primary drug fails to produce satisfactory therapeutic outcomes (*40*, *41*). Lithium is one such drug that is used for antidepressant augmentation (*9*, *41–43*). The aforementioned observation at cellular and individual levels implies that combined drug treatment targeting TCF/LEF and REST increases the likelihood of having a therapeutic effect. To test this hypothesis, we administrated mice with sertraline and lithium either in combination or separately, and assessed their effects on the behavioral phenotypes after chronic social defeat stress (Fig. 5H). We observed that the mice treated with both sertraline and lithium together showed an increased SI-rate compared to mice treated with sertraline alone (Fig. 5H). Assessment of depressive state changes in each mouse revealed that both lithium and sertraline exhibit therapeutic effects in the early week after the stressed experience (days 1–7). However, greater improvement was observed in the later week (days 8–15) with co-administration of both drugs, compared to the administration of single drugs (Fig. 5I). Collectively, these results imply that the behavioral phenotype related to chronic stress undergoes alterations due to the changes in the activity of various TFs, with the extent of influence differing among individuals; thus, applying a combination of pharmacological agents is likely to augment the therapeutic effects against depression.

## Discussion

TFs establish and regulate the transcriptome of cells. Therefore, chronic alterations in the activity of certain TFs in the brain have the potential to profoundly influence the behavioral tendencies of animals. In this study, we applied a recently developed method to evaluate TF-activities *in vivo* and discovered that TCF/LEF and REST, which displayed heightened activity in the brains of depressed mice, play a role in the recovery phase from depression. Moreover, we found lithium and sertraline, two neuropsychiatric drugs that are already prescribed, can influence the gene transcription activity of TCF/LEF and REST. These findings suggest that, notwithstanding their primary pharmacological actions, these drugs could affect the gene regulation activities of TCF/LEF and REST, thereby impacting the therapeutic outcomes.

The pathophysiology of depression remains incompletely understood, primarily due to our limited comprehension of the disease’s pathogenic heterogeneity (*14*). Our observations suggests that previously unrecognized targets of antidepressants may affect the clinical outcomes of those therapeutics. In chronically stressed mice, we observed individual differences in the activity of TCF/LEF and REST in their brains (Fig. 5A). Based on these observations, we propose that depression and the recovery from it arise from multiple mechanisms, including those related to TCF/LEF and REST. The utilization of multiple agents affecting distinct mechanisms enhances the likelihood of therapeutic success, as reflected in the switching or augmentation effect observed in medication against depression (*40–43*). Our approach, which involves analyzing the TFAP of brains, aligns well with identifying the multiple deficits regarding the function of TFs, and provide a direct means to develop therapeutic strategies against them. This will pave the path for research aiming to decipher the molecular origin of chronic disorders and foster the development of preventative or therapeutic treatments for neurological diseases. Further investigations, encompassing the acquisition of TFAPs from various neurological models and assessing how medication affects them, will substantially contribute to establishing more efficient therapeutics.

## Acknowledgments

We would like to thank N. Osumi and H. Takeuchi for their comments on the manuscript, H. Abe, A. Omori, and all other members of Abe Laboratory at Tohoku University for their help and fruitful suggestions.

## Author Contributions

Conceptualization: KA; Methodology: KA and HY; Investigation: HY, SA, RO, YG, and KA; Visualization: HY; Funding acquisition: KA, YG and HY; Project administration: KA; Writing – original draft: KA and HY; Writing – review & editing: KA and HY.

## Data and materials availability

Data and materials are provided upon reasonable request for the corresponding author.

## Materials and Methods

### Animals

Mice (*Mus musculus*, Slc: ICR) obtained through SLC Japan and bred in our facility were used. Mice in this study were maintained in clear plastic cages in a temperature- and humidity-controlled room with a 12-h light/dark cycle (light on at 8:00 a.m. and off at 8:00 p.m.) and could access standard food and water *ad libitum*. The care and experimental manipulation of animals used in this study were reviewed and approved by the Institutional Animal Care and Use Committee of Tohoku University. All experiments and maintenance were performed following relevant guidelines and regulations.

### Cell cultures and viral vectors

Cortical and thalamic neurons were obtained from mice (Slc: ICR) embryos at embryonic day 15 (E15) and E12, respectively. The cortex or thalamus was dissected and treated with 0.05 % trypsin (Fujifilm-Wako) at 37℃ following mechanical dissociation to yield single cells and then plated onto poly-L-lysine (SIGMA, #P2336) coated tissue-culture plates (IWAKI, #3820-024). The cultured neurons were maintained in Neurobasal Medium (Thermo Fisher, #21103049) supplemented with B27 Plus supplement (1:50, Thermo Fisher, #A3582801), Gultamax-I (1:100, Thermo Fisher, #35050061), and penicillin-streptomycin (1:100, Fujifilm-Wako) at 37°C under 5% CO_2_. Those neurons were transfected with viral vectors at 3 days *in vitro* (div). All of the viral vectors in this study were generated in-house, as described before (*20*). Lentivirus vectors (LV) were used for TF-activity-reporter expression, and adeno-associated viral vectors (AAV) of serotype 2/9 were used for knocking down *Lef1*, REST4 expression, and expressing the TF-activity-reporter for single-cell analysis. For CRISPR-mediated knockdown of *Lef1*, we used gRNA sequence of 5′-CAGCGACCCGTACATGTCAA-3′ and 5′-ACGGAGGCCTGTACAACAAG-3′ targeted against the open reading frame (ORF) of *Mus musculus Lef1* gene. The double-strand DNAs containing the gRNA sequence were synthesized and subcloned into the MluI/SpeI site of pAAV-FullH1TO-SaCa9sgRNAi(CREB)-CMV-TetR-2A-EGFP-KASH-WPRE-shortPA (Addgene, #113702) (*44*), which was designed to express gRNA under H1 promoter depending on doxycycline (Dox). The recombinant AAV to express SaCas9 was created using plasmid pAAV-EFS-SaCas9-P2A-HAFLAGHA-KASH-pA (Addgene, #113688). For constructs for REST4 expression, *Mus musculus Rest* transcript variant X2 mRNA sequence (XM_036164926) was synthesized with a 3× flag-tag sequence in their N-terminal. The synthesized DNA fragment was subcloned into pAAV-Syn1-Cre, an AAV expression vector with a human Synapsin promoter and WPRE-pA sequence (*20*). The LV-based TF-activity-reporters can not be utilized for RNA-seq analysis because the mRNA of its reporters and references lack poly-adenyl (pA) tails and thus can not be read by oligo-dT mediated sequence library preparation methods. Therefore, we transferred these constructs to AAV to allow the addition of pA tails to the references and reporters for single-cell RNA-sequence. To create such constructs, we cloned the LV-based reporter constructs into AAV vectors allowing bi-promoter expression of the reporter gene and reference gene by a single vector. To reduce the size of constructs, we cloned only the N-terminal region (234 bp, srtRep#5) of reporter Rep#5 and affixed to its 3’ region a unique molecular identifier (UMI) sequence (5’-CATGTTGA-3’) and a bovine growth hormone polyadenylation signal (bGHpA) sequence (5’-CTGTGCCTTCTAGTTGCCAGCCATCTGTTGTTTGCCCCTCCCCCGTGCCTTCCTTGACCCTGGAAGGTGCCACTCCCACTGTCCTTTCCTAATAAAATGAGGAAATTGCATCGCATTGTCTGAGTAGGTGTCATTCTATTCTGGGGGGTGGGGTGGGGCAGGACAGCAAGGGGGAGGATTGGGAAGAGAATAGCAGGCATTGGGGA-3’). These constructs were replaced to original reporter gene of LV-REST/NRSF-reporter (reporter gene, Rep#5; reference gene, Ref#5) (*20*). In addition, the reference gene, Ref#5, was replaced with a tagRFP gene affixed with a UMI sequence (5’-CTGCAGTA-3’) and an SV40 early polyadenylation signal sequence (5’-TCGGTCCAGTGAAAAAAATGCTTTATTTGTGAAATTTGTGATGCTATTGCTTTATTTGTAACCATTATAAGCTGCAATAAATCGAACTAGTATC-3’) to aid a visual dissection of transfected cells under fluorescent microscopes. This resulted in pAAV-REST-srtRef#5-Pgk-tagRFP. Similarly, AAV vector for TCF/LEF reporter was created from LV-TCF/LEF-reporter (reporter gene, Rep#3; reference gene, Ref#3) (*20*). The reference gene was replaced with mCherry gene affixed with UMI sequence (5’-GACTCTAT-3’) and an SV40pA sequence, and the reporter gene was replaced by N-terminal region of Rep#3 (102 bp, srtRep#3) affixed with UMI sequence (5’-CCGTATAT-3’) and bGHpA sequence, resulting in pAAV-TCF/LEF-srtRep#3-Pgk-mCherry. Plasmid used for creating AAVs for single-cell TF-activity measurement will be distributed through Addgene.

### Drug treatment

For *in vitro* drug treatment, cultured neurons were treated with drugs starting at 14 div. The cells were treated with sertraline hydrochloride (20 mM, TCI) or lithium chloride (20 mM, Nakarai-tesque) for 14–22 hours, followed by a stimulation with bicuculine methiodide (30 μM, WAKO) and 4-aminopyridine (100 μM, SIGMA) for 2 hours. Sertraline, doxycycline hyclate, and lithium chloride were applied to the medium at a final concentration of 2.5 μM, 1 μg/mL, and 50 mM, respectively. Sertraline was dissolved in dimethyl sulfoxide (DMSO, TCI), with the final concentration of DMSO being < 0.01%. For *in vivo* drug administration, drugs were diluted in drinking water and given *ad libitum*. The sertraline dissolved in DMSO was diluted to 150 mg/L with water. Either doxycycline hyclate (TCI) or doxycycline hydrochloride n-hydrate (Fujifilm-Wako) and lithium carbonate (Nakarai-tesque) were diluted to 1mg/mL and 600 mg/L, respectively.

### Surgical procedures

Mice expressing TF-activity-reporter constructs in their brains were generated as described before (*24*, *45*). Briefly, lentiviral vectors harboring the TF-activity-reporters against 6 TFs were injected into the ventricle of E15 embryos *in utero*. Mice were raised to 8 weeks when they were subjected to behavioral experiments. For injecting viral vectors into adult mice, we anesthetized mice over 8 weeks of age with the medetomidine-midazolam-butorphanol mixture (medetomidine 30 μg/mL, All Japan Pharma; midazolam 30 μg/mL, Astellas Parma; butorphanol tartrate 500 μg/mL, Meiji Seika Pharma; NaCl 118 mM; 500 μL per mouse). For knocking down the endogenous expression of *Lef1* in PFC, AAV2/9-TRE-gRNA(Lef1)-CMV-rtTA-T2A-EGFP produced from the plasmid pAAV-FullH1TO-SaCa9sgRNAi(Lef1)-CMV-TetR-2A-EGFP-KASH-WPRE-shortPA, and AAV2/9-EF1a-SaCas9-P2A-HA produced from the plasmid pAAV-EFS-SaCas9-P2A-HAFLAGHA-KASH-pA (Addgene #113688) (*46*), total ∼ 0.5 μL for each locus, were bilaterally injected at the coordinate from the bregma: anterior, +1.85 mm; lateral, ±0.35 mm; ventral, –1.7 mm of them. The same coordinates were used for virus injection for single-cell TF-activity analysis.

### Induction of depression

#### Social defeat stress

For applying social defeat stress, male mice over 8 weeks were subjected to 10 consecutive days of social defeat. Since our study used an ICR mouse for both stressor (aggressor) and defeated mice, an individual with high aggression was selected prior to the experiment and used as the stressor. For screening aggressive individuals, adult male mice aged 8 > weeks were socially isolated for more than 4 weeks, and their aggressiveness was measured regarding the frequency and severity of aggressive attacks when they were confronted with an unfamiliar male mouse. We used a blue animal marker (Asone) on the backs of aggressor mice to aid visual distinguishment of aggressive individuals. Those aggressors were used to provide social defeat stress. Before the beginning of the social defeat, the subject mice were individually housed for a week to eliminate the influence of social hierarchy established during group housing. To apply social defeat stress, we used a cage of 23 × 34 × 14 cm (CLEA Japan), divided into two compartments with transparent plexiglass with 6 holes of 1 cm diameter. The subject mouse was transferred into the aggressor’s compartment for 5 min, where it experienced physical attacks. Subsequently, the subject was moved into the opposite compartment and cohabited with the aggressor for the following 24 hours to induce psychological stress. The next day, the procedure was repeated with a different aggressor in the aggressor’s cage. This processes were repeated for 10 days with a different aggressor each day. The control mice were housed in the same type of separated cage and cohabited with the non-stressed control mice for 10 days with transferring and cohabiting to another non-stressed mouse daily.

#### Maternal separation stress

For applying maternal separation stress, mice were subjected to stress during postnatal days 2 to 15 and then raised to 8 weeks. Juvenile male mice at postnatal day 2 were randomly allocated to control or separated groups. The mice in the separated group were removed from their dams and kept in a cage maintained at 32℃ and 50–60% humidity from 4:00 p.m. to 8:00 p.m. every day from postnatal day 2 to 15. The control mice were left together with their dams. To distinguish the mice in each group, we marked the backs of both mice with animal markers. After postnatal day 16, the male mice of both groups were housed together in the same cage until they were 8 weeks old.

### Behavioral analysis

Mice were raised to 8 weeks when they were subjected to behavioral experiments. Behavioral analysis was performed in a soundproof booth equipped with a USB camera on its ceiling. Throughout the experiment, their behaviors were recorded with the camera, and their movements were tracked and analyzed using the ANY-maze software (Soldering Inc.). To acclimate them to the environment, mice with their home cage were moved to the experiment booth 30 min before the start of tests. We used a 40 × 40 × 40 cm gray plexiglass chamber for social interaction and open field tests. In the social interaction test, a display cage (circular column of 6 cm diameter, made from transparent plexiglass rods) was placed adjacent to the midpoint of a side of the chamber, and an interaction zone with a radius of 11 cm was set around it. Experimental mice were placed on the opposite side of the interaction zone and allowed to explore freely for 150 s. After the mouse was brought back to their home cage, an unfamiliar mouse was put in the display cage. Subsequently, the experimental mouse was allowed to explore again for another 150 s. Social interaction rate (SI-rate) was defined by calculating the ratio of the time spent in the interaction zone when the display cage was empty (Mouse (−)) to those when a mouse was placed in the display cage (Mouse (+)). For time-course analysis of SI-rate change, the SI-rates were normalized for the average value of the control mice at each day point. In the open field test, we recorded the trajectory of the mouse during 10 min of free exploration in that chamber. We measured the time spent in the center zone and the number of entries into the center zone. The center zone was defined as a region excluding 6 cm from the outer frame. The brightness at the center zone was 50 lux, and the rest was 30 lux. We evaluated the anxiety-like behaviors of mice with total and average per entry time spent in the center zone. In some mice, the social interaction test was conducted the day after or before social defeat, and the same individuals were used to analyze changes in sociality over time. The open field test was conducted two days after repeated social defeat or social isolation.

### Measurement of the Transcription factor activity

The endogenous activities of multiple transcription factors in the brains of mice expressing TF-activity-reporters were measured as previously described (*20*, *24*). Briefly, the TF-activity was calculated from the ratio of the expressed mRNA amount of the reporter gene and the corresponding reference gene derived from a single TF-activity-reporter construct. The reference genes are constantly expressed by a phosphoglycerate kinase (PGK) promoter, whereas the expression of reporter genes is induced by the minimal promoter with TF binding sites (TFBSs) against which the transcription factor of interest interacts. By changing the combination of a TFBS and the corresponding reference and reporter gene sets, we can measure the activities of various transcription factors from individual samples (*20*). After being subjected to experiments, the mouse brain was placed in ice-cold PBS, and then the anterior part of the cortex, where PFC was included, was rapidly dissected. Total RNAs were purified from the specimen by thiocyanate-phenol-chloroform extraction method converted to cDNAs using ReverTra Ace qPCR RT Master Mix with gDNA Remover (Toyobo, #FSQ-301). The expression of reporter and reference genes were analyzed by quantitative PCR using GoTaq-qPCR Master Mix (Promega, #A6002), with the real-time PCR system (Light Cycler 480, Roche; or CFX384, BioRad). The number of such transgenes was quantified by absolute quantification methods against standard plasmids of pre-quantified concentration. In this study, ‘TF-activity’ refers to the change of mRNA levels of the reporter gene relative to the mRNA levels expressed by reference gene from the TF-activity reporters. Certain TFs, including REST/NRSF, act as transcription suppressors depending on the cell context. However, we define increased REST-activity as an elevation in the reporter-to-reference value ratio. To measure the change in TF-activity across different conditions, sibling mice expressing the same reporter virus were randomly assigned to control and experimental groups. The activities of TFs were normalized by the average activity of that TF in control mice for each experiment. We analyzed a total of 30 TF-activity-reporters by varying the mixture of viruses to create the reporter mice. In each experiment, up to six viruses were used. These data were integrated to create a transcription factor activity profile (TFAP).

### TFAP comparison analysis

The difference in TFAPs, indicated by the Euclidean distance, represents the total distance between two plots at each vertex. To further evaluate the differences in TFAPs between groups of mice, we conducted a simulation-based analysis with acquired TF-activity data. Our simulation utilizes three steps: standardization, obtaining a value randomly, and generating simulation distribution. As each TF-activity-reporter has a different number of TFBS on their constructs and the influence of TF on their expression differs to TFs, the amplitude of measured TF-activity cannot be compared among the different TFs without correction. To treat each TF-activity-reporters equivalently, TF-activity values were corrected by subtracting the mean of and dividing by the variance of the values of the corresponding TF in the control group. Consequently, the mean and variance of TF-activity of the control group become 0 and 1, respectively. Next, the processes of randomly selecting one of the corrected values from all TFs in each experimental group and calculating those means were repeated 100 times. In this simulation, the distribution of the calculated means shows normally distributed by the central limit theorem. While the center of the distribution in the control group approximates 0, that of the experimental group having different TFAP could show values different from 0.

### Multivariate cosine similarity analysis

The analysis to identify TFs with shared change in their activity was conducted based on the cosine similarity of vectors, which comprised multiple combinations of TF-activities. Multidimensional vectors were created with changes in TF-activity (values of TFAP vertex) as components. The cosine of the angle included between and the product of the magnitude of the two vectors, whose components represented the activity change of the same TFs, were calculated for all TF-combinations. Briefly, a larger deviation in TF-activity from the control group corresponded to a larger vector size. Furthermore, a higher similarity in the pattern of TF-activity change between the two groups resulted in a higher cosine similarity. In the graphs where these values are plotted, k-means clustering was performed (k = 21; representing the number of overlapped TFs in both stressed conditions), and the dense clusters of TF-combinations in the upper-right clusters were identified through analysis using the hypergeometric distribution.

### Immunohistochemistry

Brains were perfused with 4% paraformaldehyde (Nakarai tesque) in PBS and dehydrated with 30% sucrose/PBS for 48 hours. The fixed brains were embedded into OCT compound (Sakura fintech), frozen at −80 °C, and sectioned to 40 µm thick using a cryostat (CM1850, Leica). The cells in the culture were fixed with 4% paraformaldehyde in PBS for 20 min. The sections and cultures were permeabilized with 0.1% Triton X-100 (Fujifilm-WAKO) in PBS for 15 min, blocked with 10% Normal Goat Serum (Biowest, #S1810-500) for 30 min, and stained by sequential incubation with primary antibodies diluted with PBS at 4°C overnight and with Alexa Fluor 488- or Alexa Fluor 555-conjugated secondary antibodies (1:450; ThermoFisher) or NeuroTrace500/525 (1:250; ThermoFisher, #N21480) for an hour. The primary antibodies used were as follows: anti-GFP (1:800; MBL, #M048-3MS), anti-HA (1:400; MBL, #561), and anti-LEF1 (1:200; SantaCruz, #sc374412). Fluorescent images were obtained using fluorescent microscopies with a 20 × objective lens, (Axio Imager 2, Zeiss) and a 4 × or 10 × objective lens, (BZ9000, Keyence). The number of cells and the intensity of staining were quantified by ImageJ (NIH ImageJ).

### RT-PCR analysis

The total RNA collected from the cortical culture at 14 div, and surgically dissected samples including raphe nucleus were reverse-transcribed using Revatra-ace qPCR RT Master mix with gDNA remover (Toyobo) according to the manufacturer’s protocol. The resultant cDNA was amplified by PrimeSTAR (Takara) with cycles of 95°C 10 s, 65°C 20 s, and 72°C 15 s using a PCR machine (T100, Biorad). PCR primer pairs used were 5’-TCCCAGCCTGAAAGCCACAG-3’ and 5’-ATGCTGCTGGGATTCTCTCCC-3’ for *Pet1*, and 5’-AGCTCCTGGAAGGTAAGCTTGG-3’ and 5’-GTGACAGCACTGCCCAAAAGTTA3’ for *Pgk1*. The amplified DNA products were electrophoresed on an agarose gel (Nippon-gene), stained with UltraPower DNA/RNA safe dye (Gellex), and photographed by a gel imaging device (Stage-2000, AMZ systems science), and analyzed by ImageJ.

### Transcriptome analysis

The same RNA samples utilized for measuring TF-activity were used in the transcriptomic analysis of bulk RNA-seq. The samples from the anterior cortex of control and stressed mice exhibiting social avoidance underwent mRNA purification, poly-A mRNA enrichment library preparation, and sequencing on Illumina NovaSeq-6000 platform (150 bp paired-end), performed by Novogene. For analysis of bulk-RNA-seq data, we performed differential gene expression analysis with FastQC (v0.11.9) to check raw reads and Salmon (v1.8.0) to align the reads to reference mouse genome (GRCm38) (*47*). Subsequently, we used DESeq2 software package (v1.36.0) from Bioconductor in R to perform gene expression analysis and the Benjamini-Hochberg method for multiple testing corrections. RRHO analysis was performed using RRHO2 package (version 1.0) in R. We evaluated TFBS enrichment on the sequence of the promoter region, which was obtained from UCSC Genome Browser on Mouse (GRCm39), with JASPAR2022 by TFBStools and JASPAR2022 software packages (v1.34.0 and 0.99.7, respectively) in R. Gene annotation was performed with “org.Mm.eg.db” software package (v3.15.0) in R. The heat map shown in Figure.5F was created based on transcripts per million (TPM). For this step, genes whose expression could not be detected in any of the samples were omitted, resulting in 13508 genes being visualized. These data were deposited in the DDBJ BioProject database with accession number PRJDB17042.

### Single nucleus RNA-seq analysis

For single-cell TF-reporter analysis, we injected AAV-based TF-activity reporters (AAV2/9-TCF/LEF-srtRep#3-Pgk-mCherry, AAV2/9-REST-srtRep#5-Pgk-tgRFP) into PFC of 8 weeks male mice. After recovery, these mice were subjected to chronic social defeat stress. The brains of three non-stressed control and three mice showing depressive phenotype after repeated social defeat stress were dissected, and frontal slices (∼ 1 mm thick) were created. Under a fluorescence stereomicroscope, the PFC where red fluorescent signals from reference gene (mCherry, tagRFP) were observed was dissected out (∼ 1 mm^3^) and stored frozen at −80 °C. From those frozen specimens, nuclei were isolated using a nuclei isolation kit (Minute Detergent-Free Nuclei Isolation Kit, Invent Biotechnologies, Inc., #NI-024) and collected DAPI-positive nuclei by a cell sorter (SH800, Sony). Collected nuclei were subjected to single-cell library preparation using Chromium NEXT GEM Single Cell 3’ HT Library Kit v3.1 (10× Genomics, #PN-1000370). The constructed libraries were sequenced on MGI DNBSEQ-T7 (150 bp paired-end) platform. Raw reads (about 1.2–1.6 billion in the control and defeated samples) were analyzed using Cell Ranger (v7.1.0, 10× Genomics). In concrete, resultant FASTQ files were aligned to the mouse genome (GRCm38) from 10× Genomics (https://support.10xgenomics.com/single-cell-gene-expression/software/release-notes/build) and to the specific sequence located in TF-activity-reporter constructs. Cell cluster analysis was performed with Seurat package (v4.3.0.1) in R. The normalized data of the control and defeated mice were integrated by “IntegrateData” functions in Seurat package. Following the data integration, we conducted dimensionality reduction, neighbor searching, and clustering. Subsequently, the data was visualized as a *t*-SNE plot. The expression of known marker genes was used to assign identities for each cluster: *SNAP25* for neurons, *SLC17a7* for excitatory neurons, *GAD2* for GABAergic neurons, *CUX2* for layer 2/3, *RORB* for layer 4, *TLE4* for layer 5/6 of the cortex. After clustering cells based on transcriptome data, we evaluated the average expression of the genes derived from TF-activity-reporters across all clusters in the control groups. Cells that exhibited a positive difference in the reporter-to-reference expression ratio in each cell of the defeated mice, compared to that ratio in the corresponding cluster of the control group, were categorized as having “high activity”. These data were deposited in the DDBJ BioProject database with accession number PRJDB17042.

### Quantifications and statistical analysis

The alpha score of 0.05 was used to reject the null hypothesis. All statistical analysis was performed using R. For behavioral experiments, we used Welch’s *t*-test, Tukey HSD-test, two-way ANOVA, two-way repeated-measures ANOVA followed by Tukey post hoc test, binomial test, and Pearson correlation coefficient. For a series of TF-activity analyses, we used Welch’s *t*-test, hypergeometric distribution analysis, and Pearson correlation coefficient. For a series of transcriptomic analyses, we used Student’s *t*-test, Benjamini-Hochberg procedure, paired-*t*-test, Wilcoxon rank sum test, a hypergeometric distribution analysis, and Pearson’s correlation coefficient. To measure the effect size for the difference in means for animal treatment following drug treatment, we used Hedge’s *g* statistic. The number of *n* represents the biological replicates (either individual mice or cell cultures). All boxplots display the median and the interquartile range.

## Supplementary Figures

**Supplementary Fig. S1.**
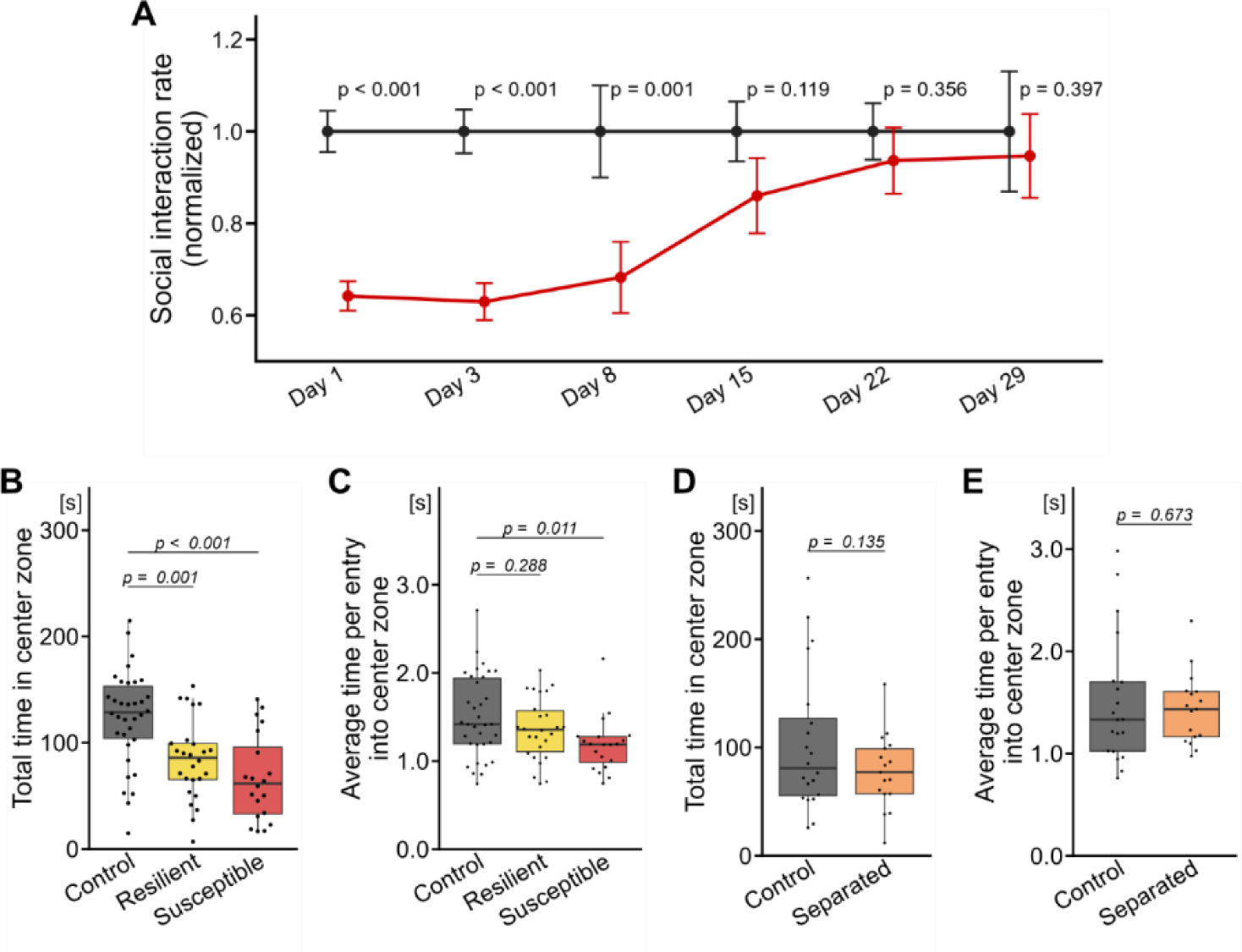
Depression-related behavior in the defeated and maternal separated mice. (**A**) Social interaction (SI)-rate change of controls and defeated mice after repeated social defeat stress. The SI-rate was normalized to control group on each day. Mean ± sem; repeated two-way ANOVA post-hoc Tukey’s test; mice per cohort; day 1, *n* = 79, 122; day 3, *n* = 79, 122; day 8, *n* = 21, 34; day 15, *n* = 18, 29; day 22, *n* = 15, 16; day 29, *n* = 15, 16. (**B**–**C**) Results of open field test on chronically defeated mice. Total (**B**) and average time spent (**C**) in the center zone. Dunnett’s test; control, *n* = 34; resilient, *n* = 26; susceptible, *n* = 20 mice. (**D**–**E**) Results of open field test on mice that had experienced maternal separation. Total (**D**) and average time spent (**E**) in the center. Welch’s *t*-test; control, *n* = 20; separated, *n* = 17 mice.

**Supplementary Fig. S2.**
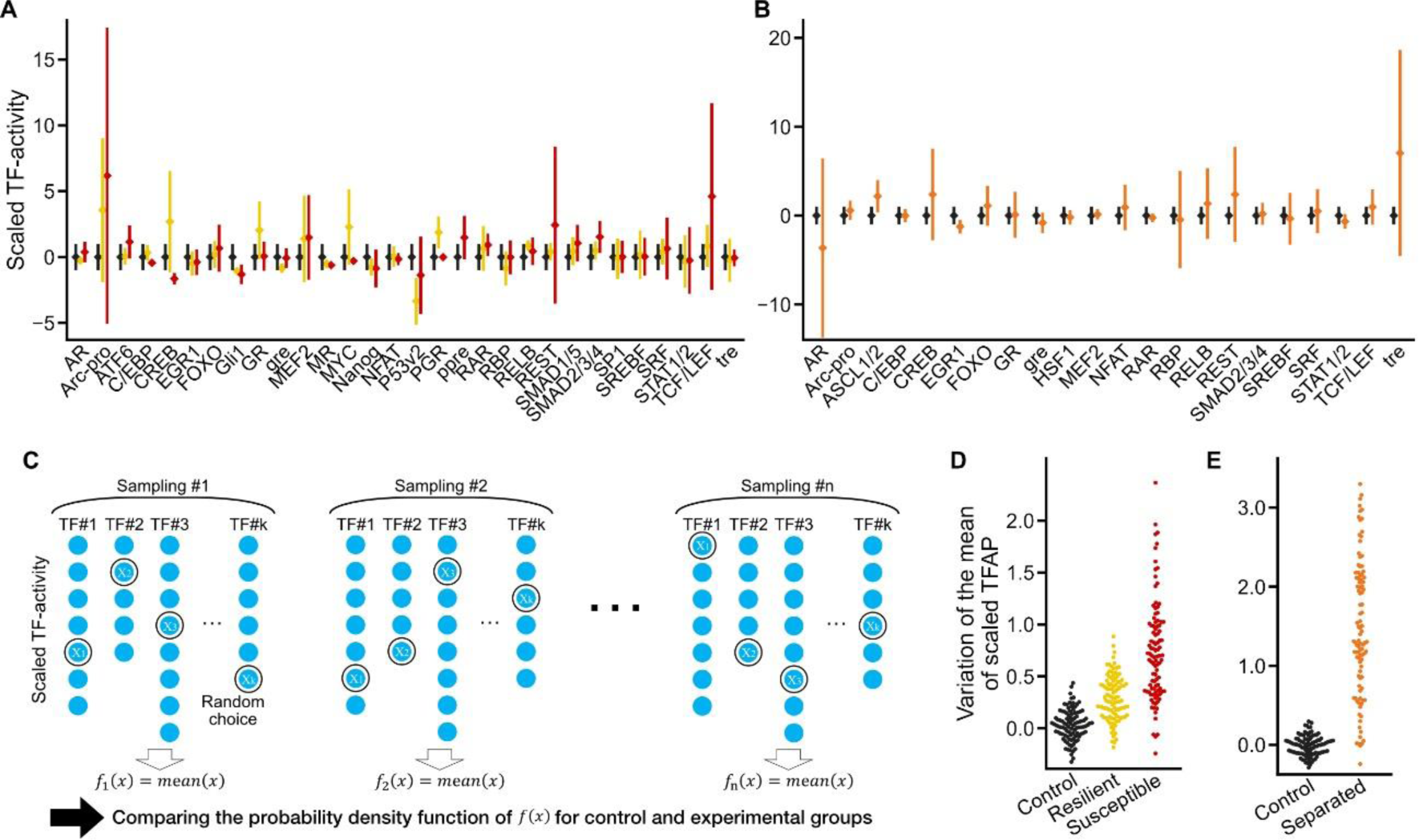
Simulation-based TFAP analysis. (**A**–**B**) Standardized TF-activity. Each measured TF-activity was corrected with the mean and standard deviation values of the control group. (**A**) Results from social defeat stressed mice, control (black), resilient (yellow), and susceptible (red). (**B**) Results from maternally separated stressed mice, control (black) and separated (orange). (**C**) Conceptual diagram of the simulation-based TFAPs analysis. In integrating the standardized TF-activities data, the challenge was that the number of data points varied for each TF. Therefore, we randomly selected one data point from each TF and integrated these. Repeating the process of calculating the average of those values 100 times resulted in a normal distribution. This procedure allows us to quantify the tendency of variance of TF-activities. (**D**–**E**) Plots showing the variation of the randomly sampled simulation results. Results from social defeat stressed mice, control (black), resilient (yellow), and susceptible (red) are shown in (**D**), and maternally separated mice, control and separated in (**E**).

**Supplementary Fig. S3.**
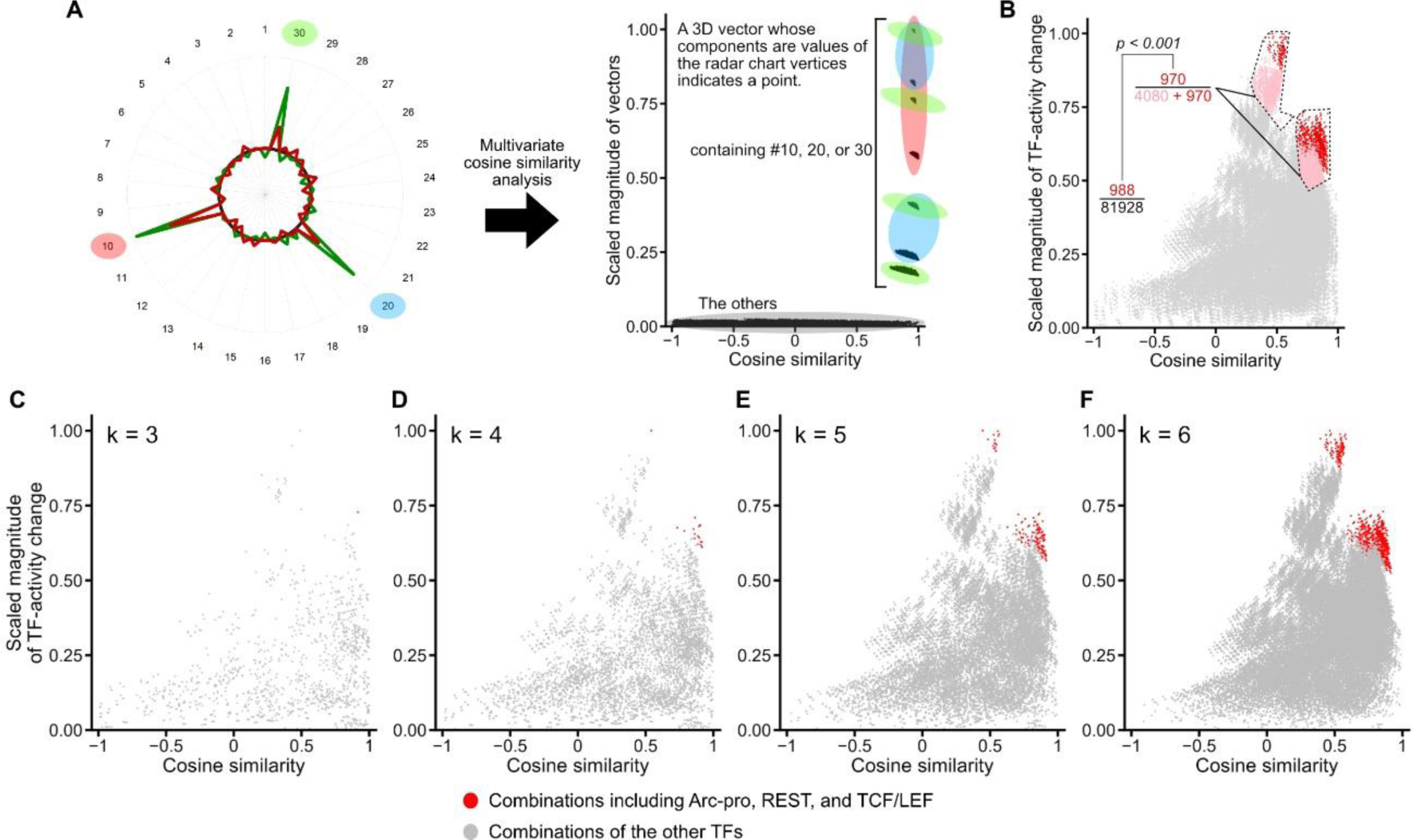
Multivariate cosine similarity analysis of the TF-activity. (**A**) Demonstration of the analysis with fictitious data. TFAPs created from fictitious data where three TFs (No. 10, 20, 30) are highly correlated among two conditions, shown as red and green TFAP (left). The result for three combinations of TFs is shown in the plot (right). The color-overlayed area includes the combination of the TFs including the TFs highlighted in the TFAP (#10, red; #20, blue; #30, green). (**B**–**F**) Results of multivariate cosine similarity the analysis of actual data obtained from susceptible mice and maternally separated mice (same as shown in Fig. 1 and 2). (**B**) The k-means clustering result shows two clusters in the upper right area, depicted in pink, with the TF combinations that include Arc-pro, REST, and TCF/LEF, highlighted in red. The inset on the left shows the ratio of combinations that include these three TF-reporters to the total number of TF combinations in the two upper right clusters or all clusters. The *p*-value is calculated based on a hypergeometric probability distribution. (**C**–**F**) The scatter plots show the results for every 3, 4, 5, and 6 combinations of TFs, respectively. These plots are overlayed in a plot shown in Fig. 2B. The combinations of TFs shown as red dots include Arc-pro, REST, and TCF/LEF.

**Supplementary Fig. S4.**
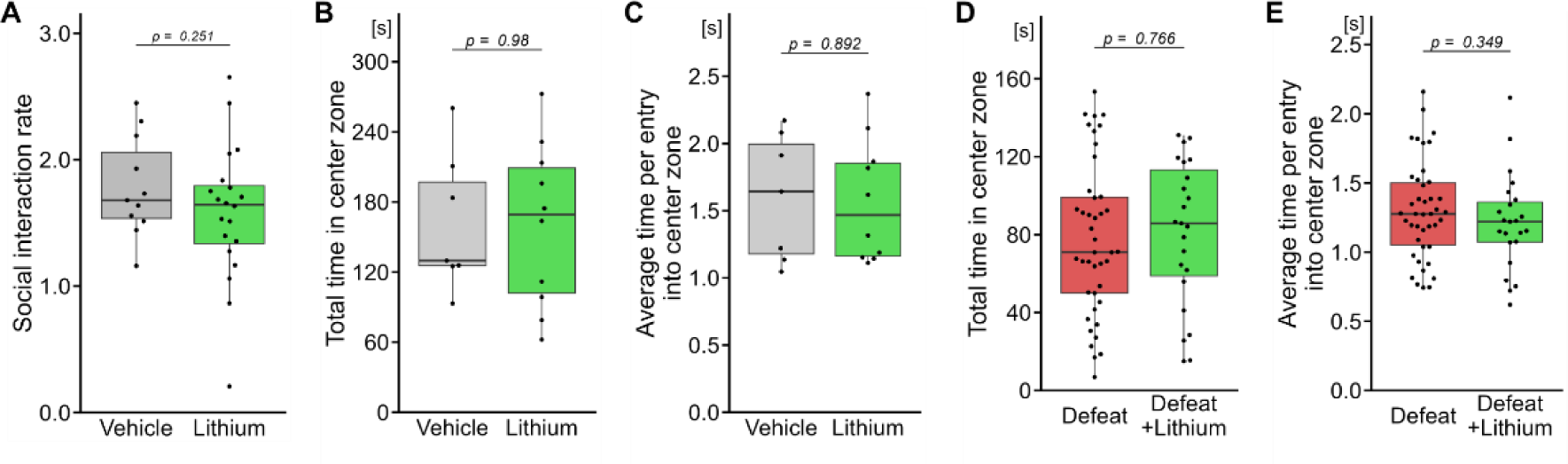
Effect of lithium administration on mice behaviors. (**A**–**C**) Behaviors of lithium-treated mice without defeat stress. (**A**) SI-rates of mice following 17 days of lithium treatment (vehicle, 11; lithium, 20 mice). (**B**–**C**) Total (**B**) and average (**C**) time spent in the center zone for those mice (vehicle, 7; lithium, 10 mice). (**D**–**E**) same as (**B**–**C**) in defeated mice with or without lithium treatment (defeat, 42; defeat + lithium, 23 mice). Welch’s *t*-test was used for the statistical analyses shown in these figures.

**Supplementary Fig. S5.**
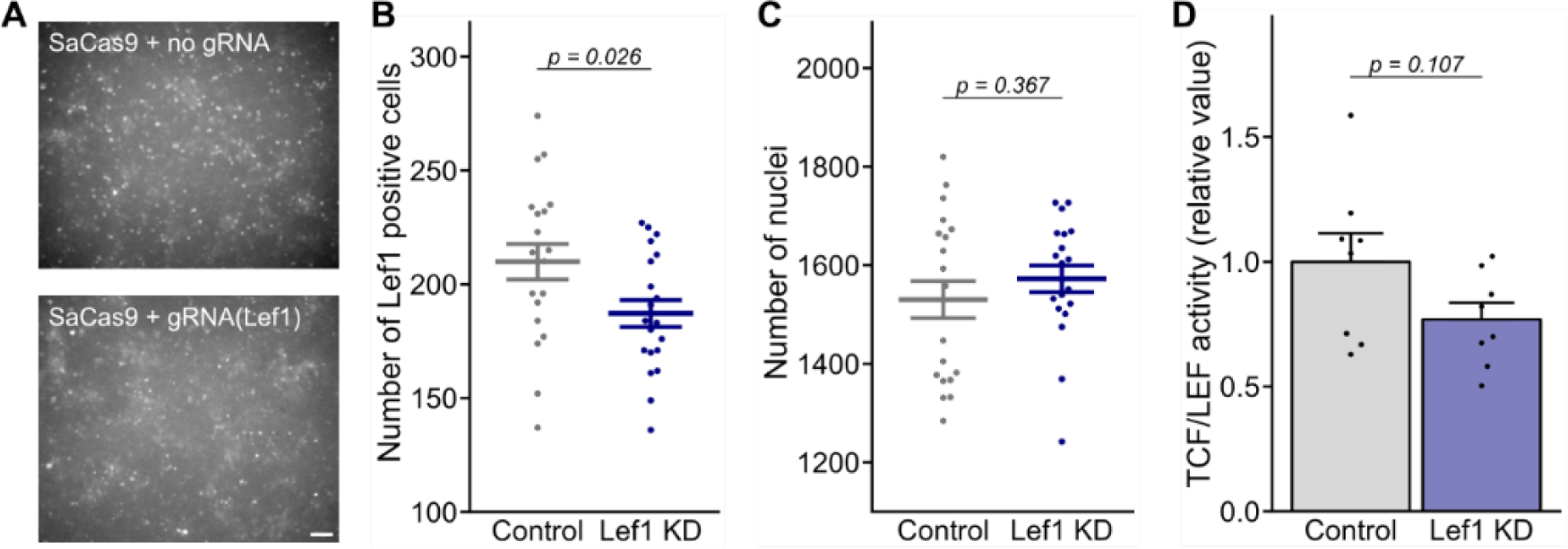
CRISPR mediated knockdown of *Lef1*. (**A**) Image of thalamic neuron culture, immunostained with anti-LEF1 antibody. The cultures were transfected with AAV to express SaCas9 without (top) or with gRNAs for *Lef1* (bottom), and treated with lithium for 24 hours from 10 days *in vitro*. Scale bar; 100 μm. (**B**–**C**) The number of cells per 50 mm^2^ positive for LEF1 (**B**) or DAPI (**C**) with or without gRNA for *Lef1* (control, 20; Lef1 KD, 20 cultures). Mean ± sem, Welch’s *t*-test. (**D**) TCF/LEF-activity of Lef1-KD cells measured by TCF/LEF-activity reporter viruses (control, 8; Lef1 KD, 8 cultures); Mean ± sem, Welch’s *t*-test.

**Supplementary Fig. S6.**
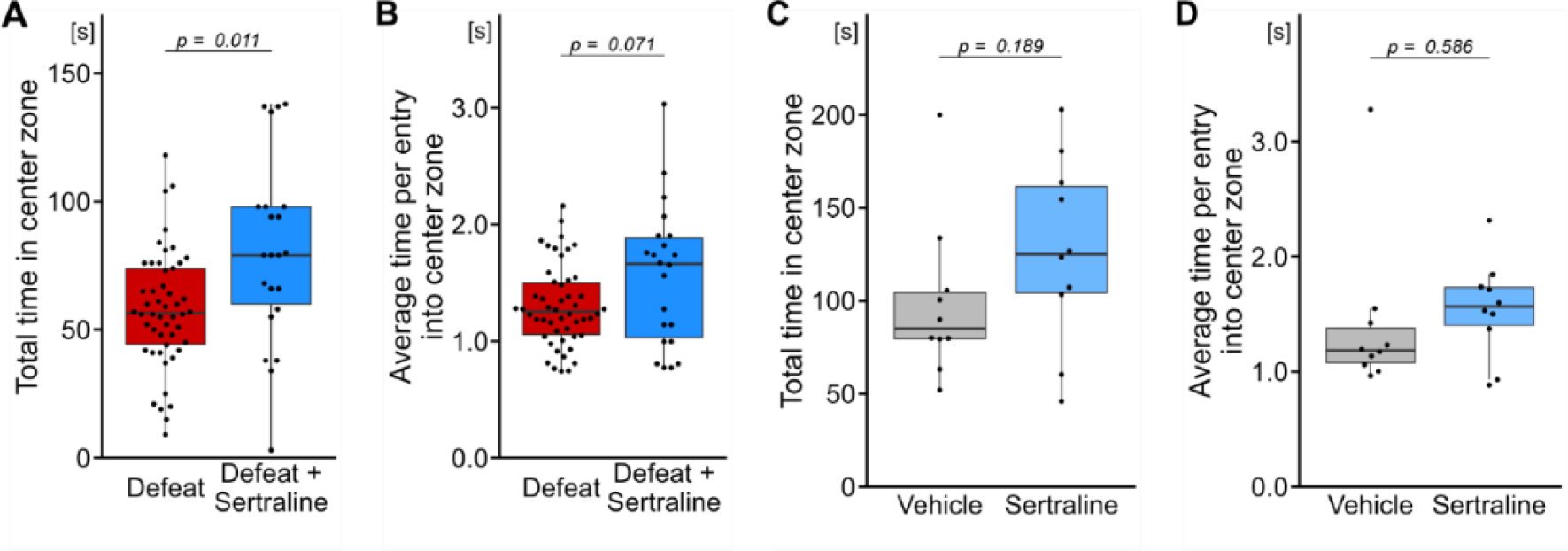
Effect of sertraline administration on mice behaviors. (**A**–**D**) Total and average time spent in the center zone in open field test. (**A**–**B**) Results from defeated mice (defeat, 20; defeat + sertraline, 22 mice). (**C**–**D**) Results from non-defeated mice (vehicle, 10; sertraline, 10 mice). Welch’s *t*-test was used for the statistical analyses shown in these figures.

**Supplementary Fig. S7.**
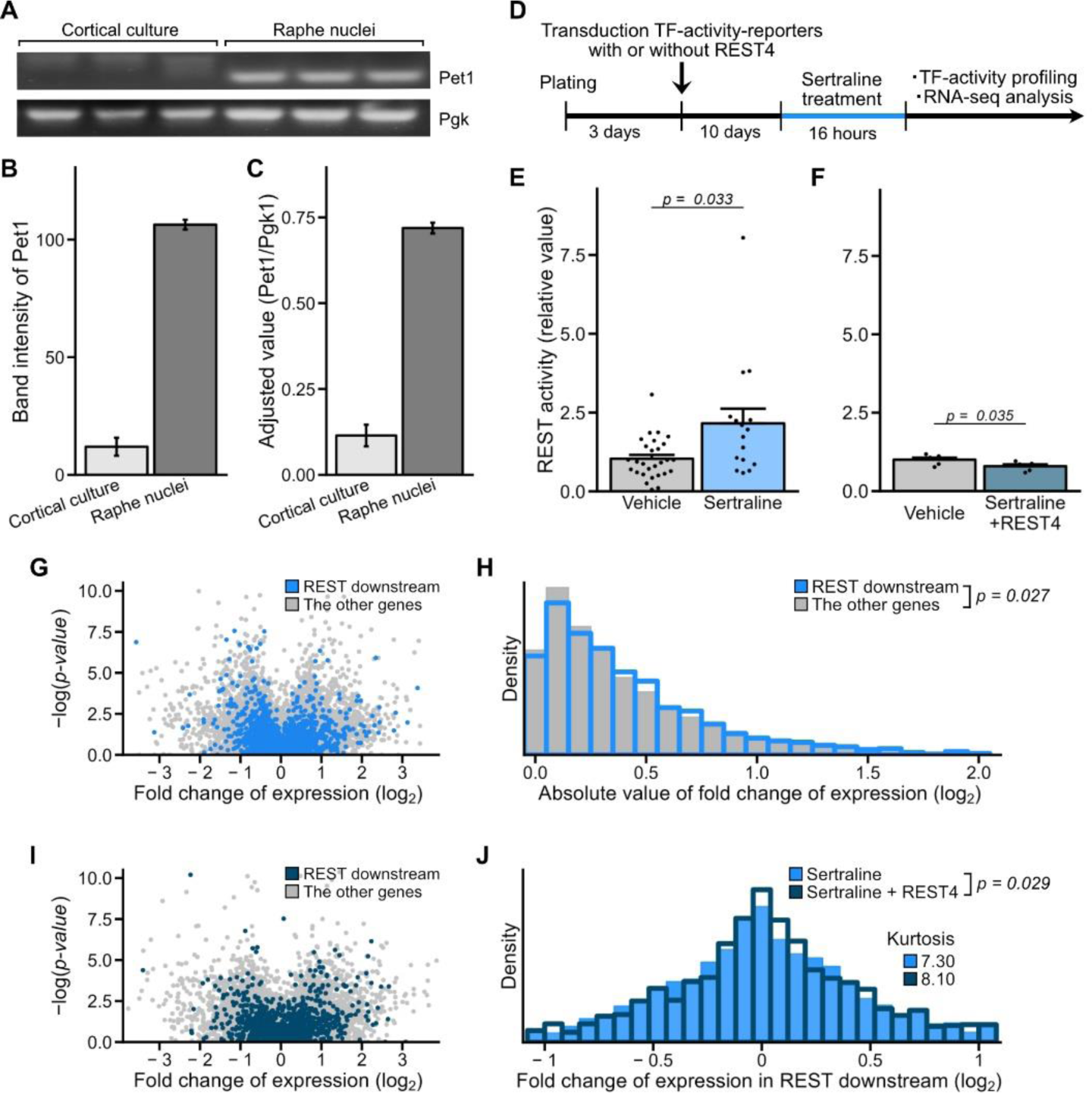
Effect of sertraline on REST-mediated gene transcription *in vitro*. (**A**) The image of gel electrophoresis showing the lack of expression of endogenous *Pet1* in cultured cortical neurons. Expression of *Pet1*, a marker for serotoninergic neurons, and *Pgk1*, a housekeeping gene, were analyzed by reverse transcription PCR (RT-PCR). (**B**– **C**) Quantification of PCR bands; mean ± sem, *n* = 3 mice or cultures. (**D**–**L**) Effects of sertraline treatment *in vitro*. (**D**) Experimental scheme of sertraline treatment *in vitro*. (**E**– **F**) REST activity in the cultured cortical neurons without (**E**) or with (**F**) REST4 expression. Welch’s *t*-test: vehicle, 28; sertraline, 16 cultures in (**E**); vehicle, 6; sertraline, 6 cultures in (**F**). (**G**) The volcano plot shows the results of RNA-seq analysis of cultured neurons; 3 biological replicates for each. Dots in light blue indicate REST downstream genes, and grey indicates the else. (**H**) Log_2_-fold-change of sertraline treated versus control neuron cultures. Wilcoxon signed-rank test; REST target, 2835; the others, 15825 genes. (**I**) same as (**G**) from cultured neurons expressing REST4. Dots in dark-blue indicate the REST downstream gene. (**J**) The histogram showing log_2_-fold-change of REST downstream genes of sertraline treated versus control neuron with or without the expression of REST4. Paired *t*-test, 2791 genes.

**Supplementary Fig. S8.**
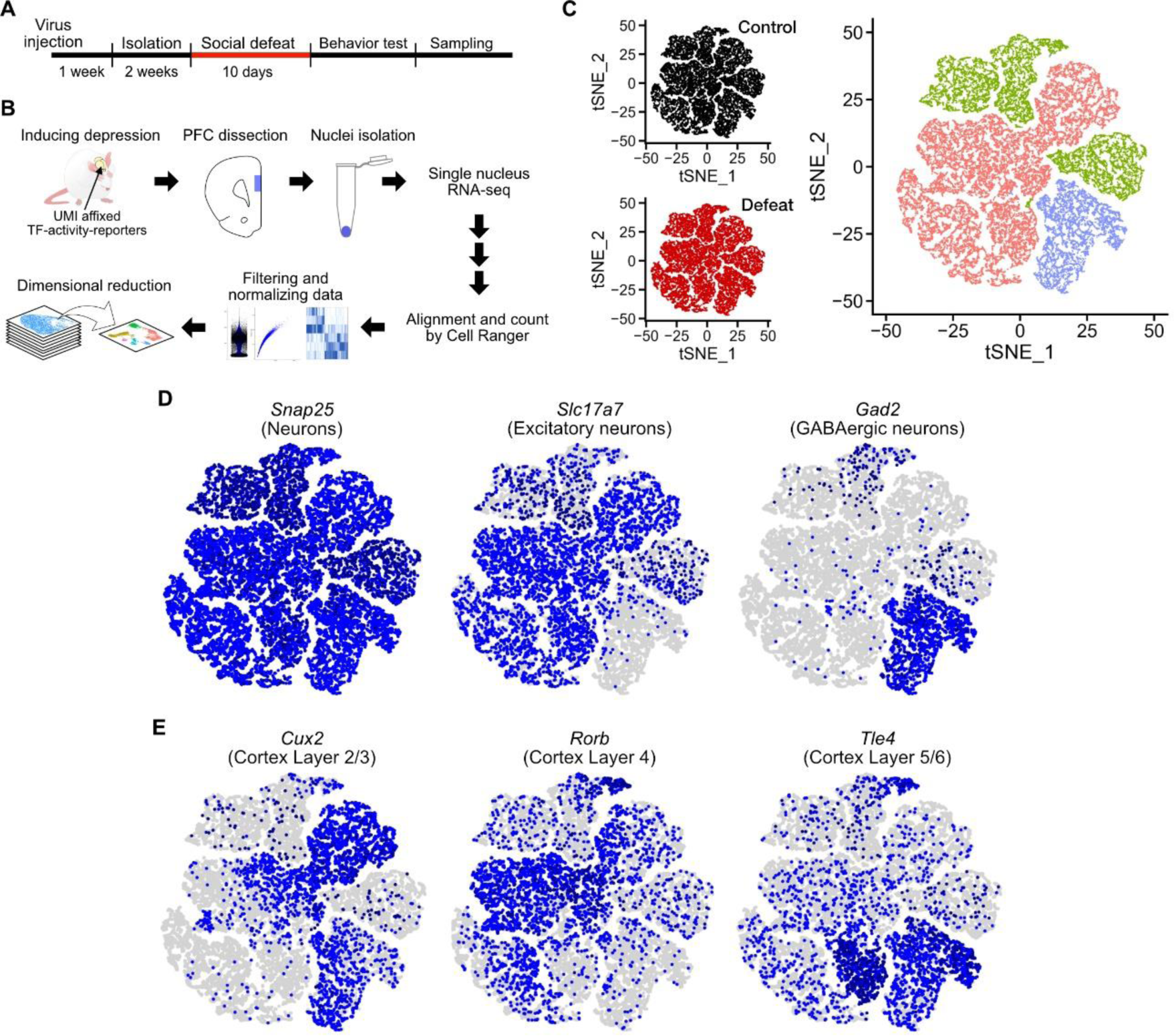
Single-cell transcriptome analysis. (**A**–**B**) The experimental scheme used for single-nucleus RNA-sequencing (**A**) and its procedure (**B**). Mice were injected with TF-reporters for REST and TCF/LEF in the PFC. (**C**) The t-SNE plots demonstrate the clustered cells using the single-cell transcriptome data. The cell distribution of cells from control (18280 cells) and defeated mice (18258 cells) (left), and the cell-identity labels (right) that are used in Figs. 5C–E. (**D**–**E)** Expression of cell-type or region-specific marker genes, with cells exhibiting high expression highlighted in blue.

